# Phage resistance accompanies reduced fitness of uropathogenic *E. coli* in the urinary environment

**DOI:** 10.1101/2021.12.02.471000

**Authors:** Jacob J. Zulk, Justin R. Clark, Samantha Ottinger, Mallory B. Ballard, Marlyd E. Mejia, Vicki Mercado-Evans, Emmaline R. Heckmann, Belkys C. Sanchez, Barbara W. Trautner, Anthony W. Maresso, Kathryn A. Patras

## Abstract

Urinary tract infections (UTIs) are among the most common infections treated worldwide each year and are primarily caused by uropathogenic *E. coli* (UPEC). Rising rates of antibiotic resistance among uropathogens have spurred consideration of alternative strategies such as bacteriophage (phage) therapy; however, phage-bacterial interactions within the urinary environment are poorly defined. Here, we assess the activity of two phages, HP3 and ES17, against clinical UPEC isolates using *in vitro* and *in vivo* models of UTI. In both bacteriologic medium and pooled human urine, we identified phage resistance arising within the first 6-8 hours of coincubation. Whole genome sequencing revealed that UPEC resistant to HP3 and ES17 harbored mutations in genes involved in lipopolysaccharide (LPS) biosynthesis. These mutations coincided with several *in vitro* phenotypes, including alterations to adherence to and invasion of human bladder epithelial HTB-9 cells, and increased biofilm formation. Interestingly, these phage-resistant UPEC demonstrated reduced growth in pooled human urine, which could be partially rescued by nutrient supplementation, and were more sensitive to several outer membrane targeting antibiotics than parental strains. Additionally, these phage-resistant UPEC were attenuated in a murine UTI model. In total, our findings suggest that while resistance to phages, such as LPS-targeted HP3 and ES17, may readily arise in the urinary environment, phage resistance is accompanied by fitness costs rendering UPEC more susceptible to host immunity or antibiotics.

**IMPORTANCE:** UTIs are one of the most common causes of outpatient antibiotic use, and rising antibiotic resistance threatens the ability to control these infections unless alternative treatments are developed. Bacteriophage (phage) therapy is gaining renewed interest, however, much like antibiotics, bacteria can readily become resistant to phage. For successful UTI treatment, we must predict how bacteria will evade killing by phage and identify the downstream consequences of phage-resistant bacterial infections. In our current study, we found that while phage-resistant mutant bacteria quickly emerged, these mutations left bacteria less capable of growing in human urine and colonizing the murine bladder. These results suggest that phage therapy poses a viable UTI treatment if phage resistance confers fitness costs for the uropathogen. These results have implications for developing cocktails of phage with multiple different bacterial targets, each of which is only evaded at the cost of bacterial fitness.

## INTRODUCTION

Urinary tract infections (UTIs) are extremely common bacterial infections, causing nearly 10 million infections in the United States alone each year(1, 2). These infections disproportionately affect women, with approximately one half of women experiencing at least one UTI during their lifetime(3). Uropathogenic *E. coli* (UPEC) are the leading cause of UTIs worldwide, causing upwards of 75% of infections each year. UTIs are one of the most common causes of antibiotic prescription in the outpatient setting(4, 5). While antibiotics are the current standard of care, the rise of antibiotic resistance among UPEC isolates threatens existing treatments for UTIs(6, 7). Current technological and economic challenges limit the development of novel antibiotics, and antibiotic resistance develops rapidly once they are introduced(8-10). Because of these challenges, several new non-antibiotic alternatives to treat UPEC UTI have been proposed(11-17). One such alternative are bacteriophages (phages); viruses that use bacteria as their natural host.

Soon after their discovery in the early 20^th^ century, phage therapy was applied to bacterial infections(18). In as early as 1928, phage therapy was applied to UTIs caused by UPEC(19). Despite this early interest, phage therapy was largely abandoned in favor of antibiotics starting in the mid-20^th^ century. As the incidence of antibiotic resistant infections increases, phage therapy is seeing a resurgence in interest(20, 21). Phage therapy holds several promises as an avenue to treat antibiotic resistant infections. Bacteriophages outnumber bacteria by an estimated 10:1 ratio worldwide(22), and phages which target human pathogens are readily isolated from environmental and human sources(23-25). Additionally, phages replicate within the bacterial host, generating a source of new phage which is limited to the duration of the pathogen presence (self-dosing). Moreover, phage may have fewer adverse or non-targeted impacts on the host microbiota(26). To date, most applications of phage therapy for UTIs, including those caused by UPEC, have been confined to compassionate care use(27-31) and have shown generally favorable results. The wide variety of pathogens and phages tested, as well as variable routes and dosage regimens limits the ability to define the clinical efficacy of phage for UTI. Clinical trials testing dosing and administration methods for UTI phage therapy are currently in the early stages of development(32, 33). The only completed randomized clinical trial conducted on UPEC UTI phage therapy described results that were non-inferior to standard-of-care antibiotics but were also non-superior to the placebo treatment(34). Factors in the urinary environment or specifically with urinary pathogens such as UPEC may impact phage efficacy and understanding phage-bacterial interactions in the urinary tract is critical to drive development of UTI phage therapies.

A key challenge to development of phage therapy for UPEC UTI is the emergence of bacterial resistance towards phage killing. Similar to the situation in which bacteria are exposed to low levels of antibiotics, bacteria can rapidly develop resistance to phage. Reported resistance mechanisms include bacterial blocking of phage adsorption through mutation of phage receptors, masking of phage targets, and producing competitive inhibitors(35, 36). Additionally, even after internalizing the phage, bacteria can resist phage infection by blocking DNA entry or degrading viral DNA once inside the cell(37). However, phage resistance has also been associated with reduction in bacterial fitness, particularly related to phage receptor modification(35, 38). These fitness costs include reduced bacterial virulence and increased susceptibility to antimicrobials(35, 39-42) or host immune clearance(43). This reduction in virulence associated with phage resistance suggests that “steering” bacteria towards this phenotype may be a viable approach for treating infections(43). Despite the current knowledge on phage resistance and its potential fitness costs, the effect of phage resistance by uropathogens in the urinary tract has not been assessed.

In this study, we evaluate the mechanisms by which UPEC evade killing by two bacteriophages that are genetically quite distinct(24) and assess the hypothesis that bacterial resistance to phage may bring associated fitness costs to the bacteria. We have found that UPEC resistance to phage killing converges on development of mutations in lipopolysaccharide (LPS) biosynthesis pathway which has detrimental effects on UPEC growth, resistance to membrane targeting antibiotics, and reduced colonization of the urinary tract *in vivo*. Together, these findings provide insight into mechanisms and associated impacts of bacterial resistance to phage in the context of uropathogenesis, and suggest phage-dependent loss of LPS reduces UPEC fitness in the urinary tract. These findings provide a critical knowledge base upon which to further develop phage therapy for UTI.

## MATERIALS AND METHODS

### Bacterial strains and mammalian cell lines

Uropathogenic *E. coli* strains UTI89 (O18:K1:H7)(44), CFT073 (O6:K2:H1; ATCC #700928)(45), DS515, and DS566 were used in isolating bacteriophage resistant bacteria. DS515 and DS566 were isolated from the urine of patients with neurogenic bladders from spinal cord injury at the Michael E. DeBakey VA Medical Center (Houston, TX). All bacterial strains were grown overnight in shaking culture at 37°C prior to experiments. Phages HP3 (accession: KY608967) and ES17 (accession: MN508615) were isolated from environmental sources and wastewater, respectively, and have been thoroughly described(24, 31). HTB-9 cells (ATCC #5367) are a human urinary bladder epithelium carcinoma cell line. Cells were grown in RPMI-1640 medium (Corning) containing 10% heat inactivated fetal bovine serum (FBS) at 37°C with 5% CO_2_ and were passaged every three to five days.

### Bacteriophage preparation

Purified stocks of phages HP3 and ES17 were prepared as previously described(46), titered, and stored in phage buffer(35) at 4°C until use.

### Human urine pool preparation

Urine samples were collected from six healthy male and female volunteers, 20-50 years-old, under approval of BCM Institutional Review Board (Protocol H-47537). Following collection, urine was warmed to 37°C and filtered using a 0.22µm filter before storage at 4°C.

### Isolation of phage-resistant bacteria

Overnight bacterial cultures were diluted in fresh LB or pooled human urine and approximately 10^8^ CFU UPEC was added to 96-well microtiter plate wells and were challenged with 10^7^ PFU phage (MOI = 0.1) in a total volume of 150µL. PBS without phage was used to assess non-infected bacterial growth. Optical density was measured every 15 minutes using a Tecan Infinite 200 plate reader (Tecan i-control version 1.7). After 18 hours, phage-treated wells with bacterial growth were streaked onto soft agar overlay plates containing 1×10^8^ PFU of phage. This process was repeated once for colonies which grew on phage top agar to isolate clonal populations. Resistance to phage was confirmed by spot assay onto lawns of UPEC.

### Purification and visualization of LPS

UPEC LPS was isolated through hot aqueous-phenol extraction following previously described methods(47). Extracted LPS samples were run on 4-12% SDS-polyacrylamide gels with 15µL of extracted sample loaded into each well. Samples were run at 120V for 1.5h and were stained using the Pro-Q Emerald 300 Lipopolysaccharide Gel Stain Kit (Molecular Probes) following manufacturer directions. Gels were visualized utilizing the ProteinSimple AlphaImager HP system and associated software.

### Generating bacterial growth curves

Overnight cultures of UPEC were diluted 1:100 in either LB or pooled human urine and added to 96-well microtiter plates. For phage challenge growth curves, bacteriophage preparations diluted in PBS were added at MOI of 0.1, 0.001, or 0.00001 to reach a final volume of 150µL. Control bacterial growth wells were treated with PBS alone and all conditions were tested in technical duplicate. Growth curves were generated using a Tecan Infinite 200 plate reader at 37°C measuring OD_600_ every 15 minutes for 18h. “Relative bacterial growth” was determined by calculating the percent OD_600_ of a given well compared to the mean of untreated wells at that same timepoint. For growth assessments of phage-resistant bacterial growth, overnight cultures were diluted 1:100 in LB media or pooled human urine and added to 96-well microtiter plates in a total volume of 100µL. Plates were grown in a shaking BioTek Cytation 5 plate reader (Gen5 software version 3.10) at 37°C measuring OD_600_ every 15 minutes for 16h.

### Biofilm assays

Bacterial biofilms were quantified as previously described with minor adaptations(48). Briefly, overnight cultures were diluted to OD_600_ 0.1 in either LB, urine, or RPMI-1640 before adding 200µL to 96-well tissue culture plates. Plates were incubated in stationary conditions at 37°C for 24 h. The following day, OD_600_ was measured to quantify overall bacterial growth. Non-adherent bacteria were removed, and washed three times with PBS. Biofilms were then dried at 55°C for 1h. Afterwards, 200µL of 0.2% crystal violet solution was added to each well and plates were incubated at room temperature for 30 minutes. Crystal violet was removed, and plates were washed five times with PBS. To release crystal violet, 200µL of an 80:20 mixture of ethanol and acetone was added to each well. The released crystal violet solution (100 µL) was transferred to a new 96-well plate and absorbance was measured at OD_595_ on a BioTek Cytation 5.

### Adherence and invasion assays

Adherence and invasion assays were performed using previously described methods(49, 50). The day prior to adherence and invasion assays, HTB-9 cells were passaged from a T-75 tissue culture flask into 24-well tissue culture plates and grown at 37°C with 5% CO_2_ overnight. Confluency was assessed the following day prior to experiment. The day of the experiment, overnight bacterial cultures were subcultured and allowed to reach mid-log phase (OD_600_ = 0.4 - 0.6). Thirty minutes prior to adding bacteria, HTB-9 cells were washed and 400µL of fresh RPMI-1640 media was added. Bacterial inoculum was prepared by diluting mid-log phase cultures 1:10 in PBS. 100µL of bacterial dilution was added to wells and plates were spun at 200 x *g* for two minutes to facilitate bacterial-cell contact. Plates were then incubated at 37°C in a 5% CO_2_ incubator. For adherence assays, after 30 minutes of incubation, cells were washed six times with PBS before the addition of 100µL of 0.025% Trypsin-EDTA. Cells were incubated at 37°C for seven minutes, after which time, 400µL of 0.025% Triton X-100 was added to each well to lyse HTB-9 cells. Wells were pipetted up and down 25 times before being diluted and plated to assess bacterial burden. For invasion assays, infected HTB-9 cells were incubated with bacteria for two hours, after which time, media was removed and replaced with 500µL of RPMI-1640 media containing 50µg/mL gentamycin. The cells were again incubated for two hours before being washed and lysed as described for adherence assays.

### Minimum inhibitory concentration (MIC) assays

MIC assays were conducted as described previously(51). Briefly, overnight cultures of bacteria were subcultured in LB media and allowed to reach mid-log phase (OD_600_ = 0.4 - 0.6) before being pelleted and resuspended 1:10 in PBS. Antibiotic stocks were diluted in LB or urine and added to the top row of 96-well plates before being diluted twofold down the plate. Bacteria were added at a 1:10 dilution to each well before being incubated at 37°C overnight. The following day, absorbance at OD_600_ was measured using a BioTek Cytation 5 plate reader. To generate a greater dynamic range to assess bacterial growth, resazurin was added at 6.75 μg/mL to each well and plates were incubated for an additional three hours at 37°C in the dark. Following this incubation, fluorescence was measured using excitation/emission of 550/600 on a BioTek Cytation 5, and MIC was determined as the lowest antibiotic concentration at which 90% growth suppression occurred.

### Animals

All animal experiments were approved by the Baylor College of Medicine (BCM) Institutional Animal Care and Use Committee (Protocol AN-8233) and were performed under accepted veterinary standards. Female C57BL/6J mice (Strain #000664) were purchased from Jackson Laboratories or from BCM vivarium stock and all experiments were conducted when mice were aged 8 to 12 weeks. Animals were allowed to eat and drink *ad libitum* throughout the duration of experiments.

### Murine UTI model experiments

An established murine UTI model was used as previously described(50, 51). Mice were anesthetized with inhaled isoflurane and approximately 1 × 10^8^ CFU of bacteria were transurethrally instilled into the bladders of mice in 50µL liquid volume. Transurethral infection was achieved by inserting polyethylene tubing (inner dimension: 0.28 mm; outer dimension: 0.61 mm) covering a 30-gauge hypodermic needle into the urethra of anesthetized mice. At 24 hours post infection, bladders and kidneys were removed from each mouse and homogenized in tubes containing 1.0mm diameter zirconia/silica beads (Biospec Products, catalog no. 11079110z) using a MagNA Lyser (Roche Diagnostics). Serial dilutions of homogenized organs were plated on LB agar and enumerated the following day.

### Sequencing and analysis of phage resistant UPEC

Bacterial DNA was isolated from overnight bacterial cultures using the E.Z.N.A. bacterial DNA kit (Omega Bio-Tek) following manufacturer instructions. Sequencing was performed by Novogene using the Illumina Platform. Reads were trimmed to Q30 and a minimum length of 50 bp using BBDuk (version 38.84). Phage-resistant isolates were compared to wild-type strains using three comparison methods: (1) reads were independently assembled *de novo* using Geneious assembler in Geneious 2022.0.1. Contigs of resistant bacteria were then compared to parental contigs using progresssiveMauve (version 2015-02-26) and disagreements were extracted(52); (2) Reads from phage-resistant bacteria were mapped to the wild-type bacteria from method 1, followed by variant analysis using Geneious 2022.0.1 variant finder; (3) Resistant bacteria were compared to parental bacteria using snippy-multi script in Snippy(53). Snippy results agreed with results from methods 1 and 2 and are presented here. For reference selection in Snippy, bacterial reads were validated through EDGE Bioinformatic (version 2.4.0) software’s phylogenetic analysis module, using RAxML and a pre-built *E. coli* SNP database(54, 55). The published UTI89 genome (accession: CP000243.1) was used as a reference. Since DS566 did not cluster with another strain closely enough for that strain to be used as a reference, it was further categorized using the Center for Genomic Epidemiology’s MLST 2.0 software (version 2.0.4; database version 2021-10-18) which predicted it belonged to ST1193(56, 57). *E. coli* MCJCHV-1 (accession: CP030111.1) was chosen as a reference for DS566. Variants that were present in both parental strains and resistant progenies were discarded. Circular diagrams were made using CGView Server and modified using Microsoft PowerPoint(58).

### Statistics

*In vitro* experiments were performed at least twice in at least technical duplicate. Mean values of independent experiments were used to represent biological replicates for statistical analyses. *In vivo* experiments were conducted at least twice independently with individual mice serving as biological replicates. Experimental data was combined prior to statistical analyses. Mann-Whitney tests were used to compare AUC measurements (Fig. 1E-F), and murine bladder colonization (Fig. 6A-B). Friedman test using Dunn’s multiple comparisons was used to assess changes in bacterial adherence and invasion (Fig. 3A-D), while two-way repeated measures ANOVA with Geisser-Greenhouse correction and Dunnett’s multiple comparisons was used for biofilm experiments (Fig. 3E-F). Statistical analyses were performed using GraphPad Prism, version 9.2.0 (GraphPad Software Inc., La Jolla, CA). *P* values of <0.05 were considered statistically significant.

**Figure 1:**
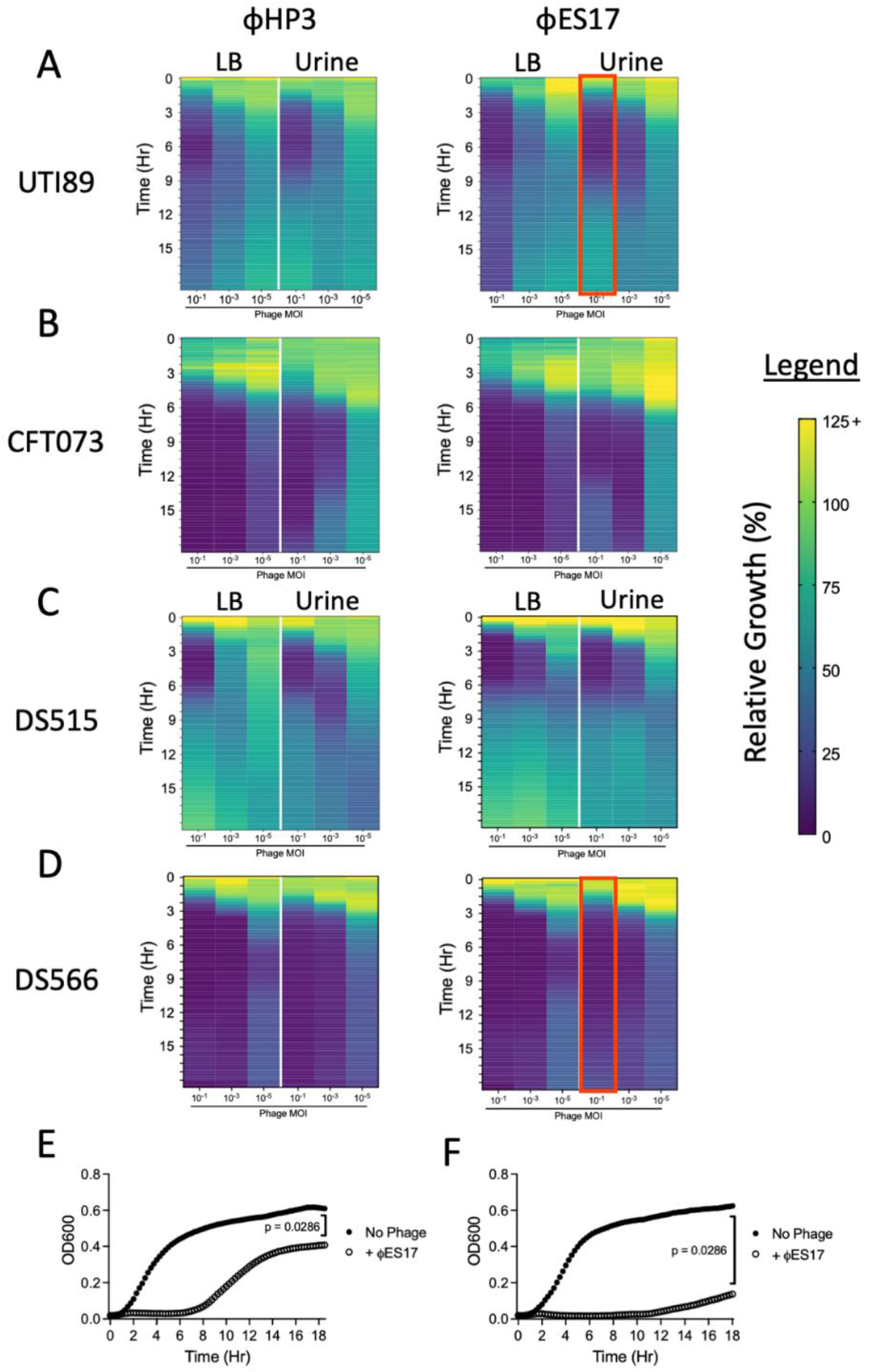
Phage-bacterial dynamics are similar in LB and pooled human urine with resistance developing in both conditions. Heatmaps of relative bacterial growth (OD_600_ of phage treated well/OD_600_ of untreated well) of bacteria challenged with HP3 (left column) or ES17 (right column) in LB media and pooled human urine during 18h of growth. Conventional UPEC strains UTI89 (**A**) and CFT073 (**B**) were used as well as recently isolated catheter associated UTI isolates DS515 (**C**) and DS566 (**D**). All bacteria were challenged at multiplicities of infection (MOI) of 10^−1^, 10^−3^, and 10^−5^. (**E**) Representative growth curve of UTI89 challenged with ES17 at MOI 10^−1^ in urine highlighted in red in (**A**). (**F**) Representative growth curve of DS566 challenged with ES17 at MOI 10^−1^ in urine highlighted in red in (**D**). All heatmaps are representative of two to three independent experiments performed in duplicate or triplicate. Area under the curve (AUC) was measured for each growth curve and analyzed by Mann-Whitney test.

## RESULTS

### UPEC resistant to phage rapidly emerge *in vitro* in both LB and human urine

Conventionally, bacteria-phage interactions are studied in bacteriologic medium, however, more recent work has begun to assess these interactions in the context of the host environment including *ex vivo* blood or *in vivo* tissues(43, 59, 60). To assess how phage activity could be influenced by the urinary environment, we compared 18h bacterial growth curves in LB medium and human urine. We selected two phages(24, 31) to test against four UPEC strains. UTI89 and CFT073 are well-characterized UPEC cystitis and pyelonephritis isolates respectively, while DS515 and DS566 are recently isolates from patients with neurogenic bladders from spinal cord injury. HP3 and ES17 are phages with demonstrated efficacy against extraintestinal pathogenic *E. coli* (ExPEC) including UPEC(24, 31, 35, 46, 61).

Approximately 10^8^ CFU of UPEC were challenged with 10^7^, 10^5^, or 10^3^ PFU of phage (MOI of 10^−1^, 10^−3^, and 10^−5^ respectively) in 96-well microtiter plates. Within the first 1-2h of phage challenge, relative growth of phage treated wells rapidly declined. Minimal growth (< 25% growth of OD_600_ in treated vs untreated wells) was observed for the first approximately 6h in the presence of the two higher MOIs for both phages (**Fig. 1A-D**). After this time, many of the cultures began to “rebound” their relative growth, representing bacterial resistance towards phage. This activity was observed most strongly in strains UTI89 (**Fig. 1A**) and DS515 (**Fig. 1C**). There were no clear differences in relative growth based on medium used (LB vs urine), although decreasing phage MOI caused less suppression of relative growth. Bacterial strains UTI89 and DS566 were selected for further investigation of the mechanisms of phage resistance due to their differential growth kinetics observed during *in vitro* phage challenge. While relative growth was quick to rebound when UTI89 was challenged with phage (**Fig. 1E**), DS566 relative growth was slow, reaching only ∼20% of no phage control growth after 18h of incubation (**Fig. 1F**).

### Rebound in bacterial growth is due to resistance to phage through mutations of LPS

To isolate phage-resistant UPEC, bacteria were passaged twice in the presence of phage in liquid culture and on agar containing phage (**Fig. 2A**). Three UTI89 isolates resistant to HP3 or ES17 and two DS566 isolates resistant to ES17 were collected for whole-genome sequencing. Sequencing revealed that all UTI89 resistant to phage, regardless of which phage, had mutations in the transcription factor rfaH (locus tag: b3842, **Fig. 2B**). DS566 mutants had mutations in galactotransferase rfaI/waaO (locus tag: b3627). DS566-2 had an additional mutation in rfaE/hldE (locus tag: b3052, **Fig. 2C-D**). Several non-LPS related mutations were also observed (**Supp. Table 1**). When LPS was isolated through hot aqueous-phenol extraction, the loss of LPS O-antigen was apparent in all phage-resistant strains (**Fig. 2E**). These results agree with LPS structure prediction with mutations in rfaH and rfaE/hldE predicted to result in truncation of the inner core of LPS while rfaI/waaO is likely to result truncation of the outer core of LPS(62) (**Fig. 2F**).

**Figure 2:**
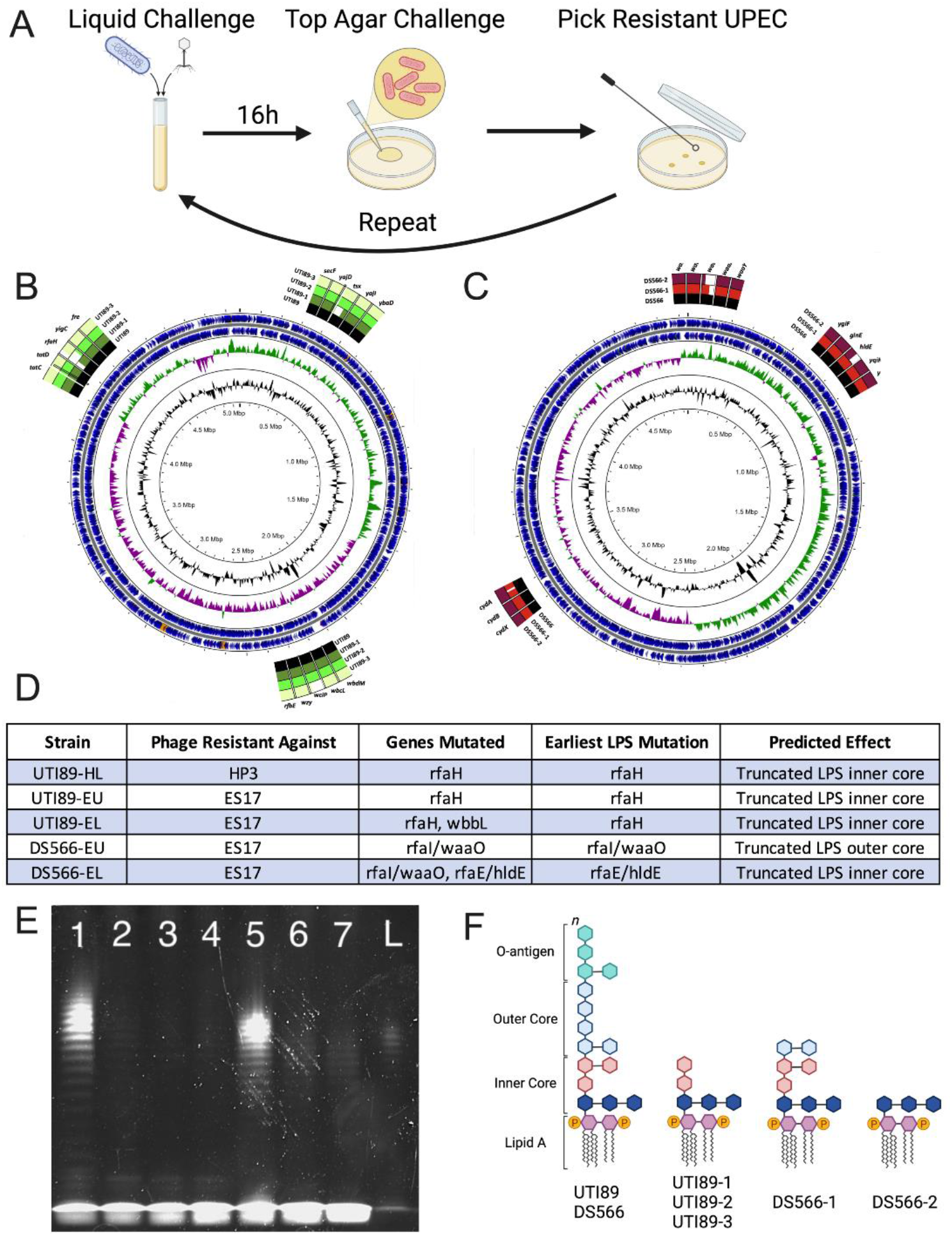
Isolation and sequencing of phage resistant bacteria indicates mutations to LPS biosynthesis confer resistance to phages HP3 and ES17. (**A**) Schematic demonstrating the selection protocol for identifying clonal phage-resistant mutant UPEC through serial phage challenges. (**B**) Diagrams depicting mutations identified in UTI89 phage-resistant mutants and (**C**) DS566 phage-resistant mutants. (**D**) Summaries of mutations identified through whole genome sequencing. (**E**) Representative SDS-page image of wild-type and phage resistant UPEC LPS isolated through hot aqueous-phenol extraction run on a 4-12% SDS-polyacrylamide gel loaded with 15µL of isolated product. Lane 1: Wild-type UTI89, lane 2: UTI89-1, lane 3: UTI89-2, lane 4: UTI89-3, lane 5: Wild-type DS566, lane 6: DS566-1, lane 7: DS566-2, L: LPS standard. Isolation and visualization of LPS was performed in two independent experiments with comparable results. (**F**) Predicted LPS structures based on LPS mutations noted in sequencing analysis. Images for panels A and F were created using BioRender software.

### LPS mutation leads to changes in biofilm formation, adherence and invasion of bladder cells, and antibiotic susceptibility

UPEC adheres to and invades the bladder epithelium to form intracellular bacterial reservoirs capable of reseeding infection(63, 64), thus, we assessed if LPS mutations confer changes in the ability of UPEC to adhere to and invade HTB-9 bladder cells. No change in adherence was seen in any of the UTI89 phage resistant mutants (**Fig. 3A**), however, both DS566 mutants had modestly increased adherence relative to the wild-type strain (**Fig. 3B**). Invasion of HTB-9 cells was increased 50 to 100-fold in two of three UTI89 mutants, UTI89-1 and UTI89-3 (**Fig. 3C**), while increased invasion was only detected in one of the DS566 mutants, DS566-1 (**Fig. 3D**).

**Figure 3:**
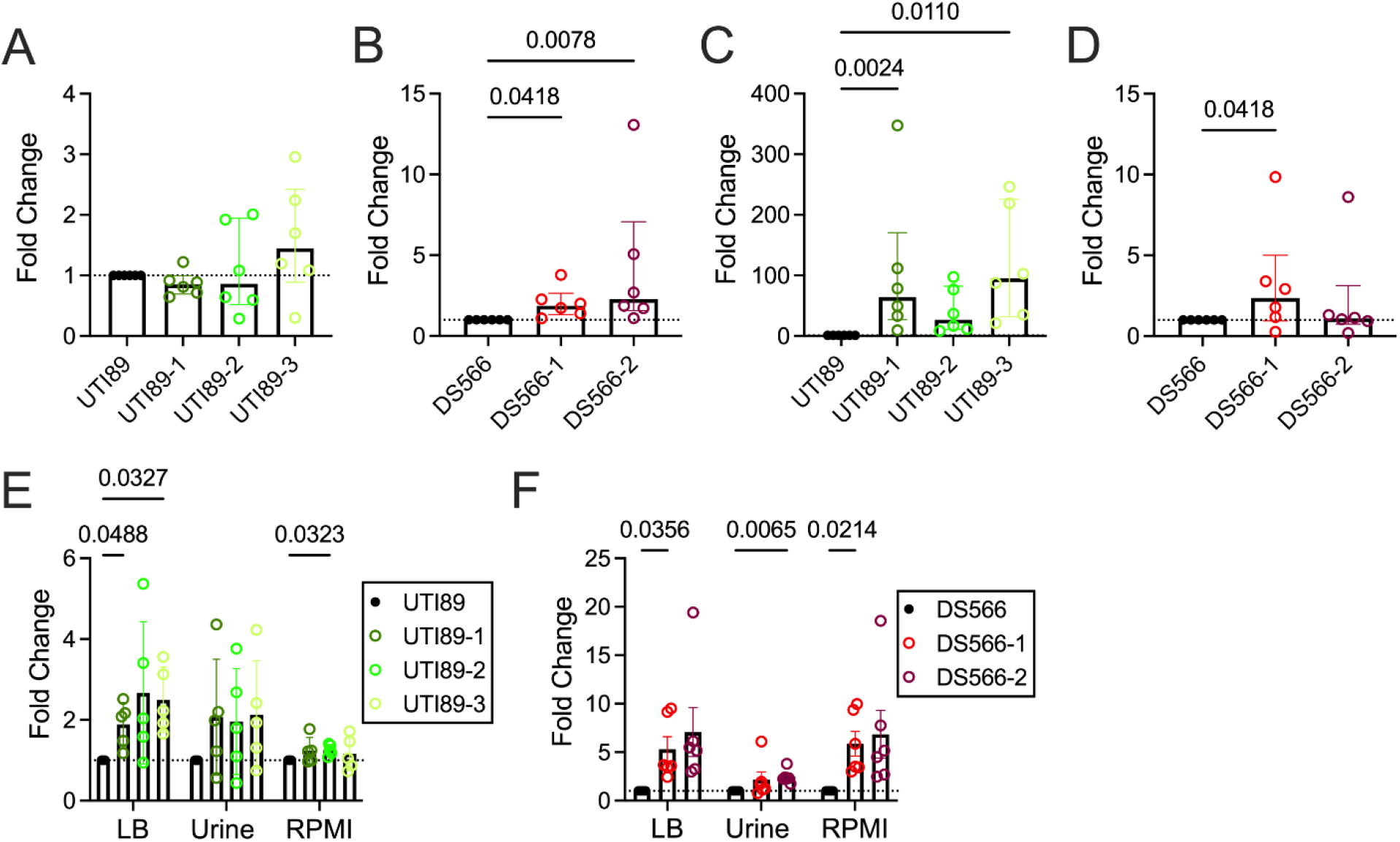
Adherence and invasion of HTB-9 cells as well as biofilm formation is altered in LPS mutant UPEC. (**A**) UTI89 and (**B**) DS566 and respective LPS mutant bacterial adherence to HTB-9 cells after 30 min of infection, MOI = 1. (**C**) HTB-9 cells were infected with UTI89 or (**D**) DS566 and their LPS mutants (MOI = 1) for two hours before being media was changed to RPMI-1640 containing 50µg/mL gentamycin to kill extracellular bacteria. After two hours of antibiotic treatment, cells were lysed, and intracellular bacteria enumerated. (**E**) Biofilm formation of UTI89 or (**F**) DS566 and their LPS mutants in LB, urine, or RPMI-1640 quantified by crystal violet uptake. All adherence, invasion, and biofilm assays were performed in five to six independent experiments. Adherence (A,B) and invasion (C,D) assays were performed in technical duplicate or triplicate while biofilm (E,F) assays were performed with six to eight technical replicates. Data was analyzed by Friedman test using Dunn’s multiple comparisons test (A-D), or two-way repeated measures ANOVA with Geisser-Greenhouse correction and Dunnett’s multiple comparisons test (E,F).

As LPS mutations are likely to change bacterial surface properties(65, 66), we evaluated the ability of these bacteria to form biofilms in LB as well as more physiologically relevant conditions of pooled human urine and RPMI-1640 medium. In general, biofilm formation by phage-resistant UPEC was increased relative to the parental strain although not all conditions achieved statistical significance as indicated in figure panels. UTI89-1 and UTI89-3 demonstrated significantly increased biofilm formation in LB medium while UTI89-2 demonstrated enhanced biofilm formation in RPMI-1640 (**Fig. 3E**). DS566-1 displayed increased biofilm formation in both LB and RPMI-1640 compared to its wild-type strain, whereas DS566-2 had increased biofilm formation in pooled human urine (**Fig. 3F**).

LPS truncation may allow antimicrobials to access the bacterial surface more easily. We tested if the antibiotics colistin (polymyxin E) and polymyxin B, which both interact with the bacterial outer membrane, had lower minimum inhibitory concentration (MIC) values for LPS-mutant UPEC compared to wild-type strains in pooled human urine. We observed a 2-fold decrease in the colistin MIC in UTI89 LPS mutants UTI89-2 and UTI89-3 (**Fig. 4A**), and a non-significant change in MIC versus polymyxin B (**Fig. 4B**). In DS566, we observed a 2-fold reduction in MIC for LPS mutant DS566-1 to both colistin and polymyxin B while DS566-2 had colistin and polymyxin B MICs that were 8 to 16-fold lower than the wild-type strain (0.1875 µg/mL and 0.09375 µg/mL, respectively) (**Fig. 4C-D**). In LB medium, differences between phage-resistant mutants and parental strains were largely absent (**Supp. Fig. 1**).

**Figure 4:**
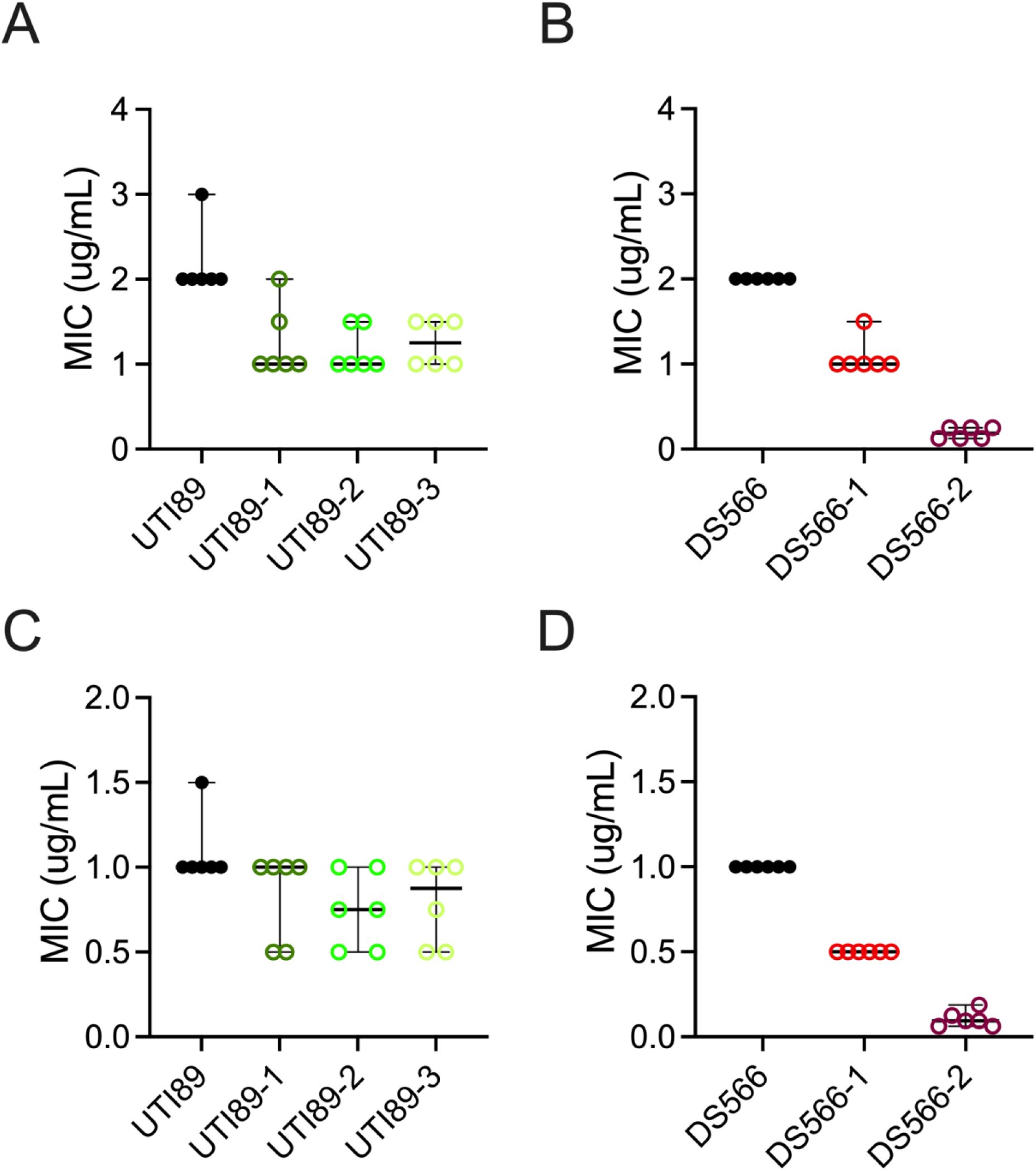
LPS mutation renders UPEC more susceptible to antibiotics that target the bacterial outer membrane. Colistin MICs of (**A**) UTI89 and (**B**) DS566 and their LPS mutants in pooled human urine. Polymyxin B MICs of (**C**) UTI89 and (**D**) DS566 and their LPS mutants in pooled human urine. Individual points are representative of independent experiments performed in duplicate. Bars represent median and 95% confidence intervals.

### LPS-mutant UPEC growth is attenuated in urine

To evaluate the impact of LPS modifications on bacterial growth, bacteria were grown in either LB medium or pooled human urine for 16h. While growth of LPS mutants was similar to the wild-type strain in LB medium (**Fig. 5A-B**), bacterial growth was severely attenuated in urine, regardless of specific LPS mutations (**Fig. 5C-D**).

**Figure 5:**
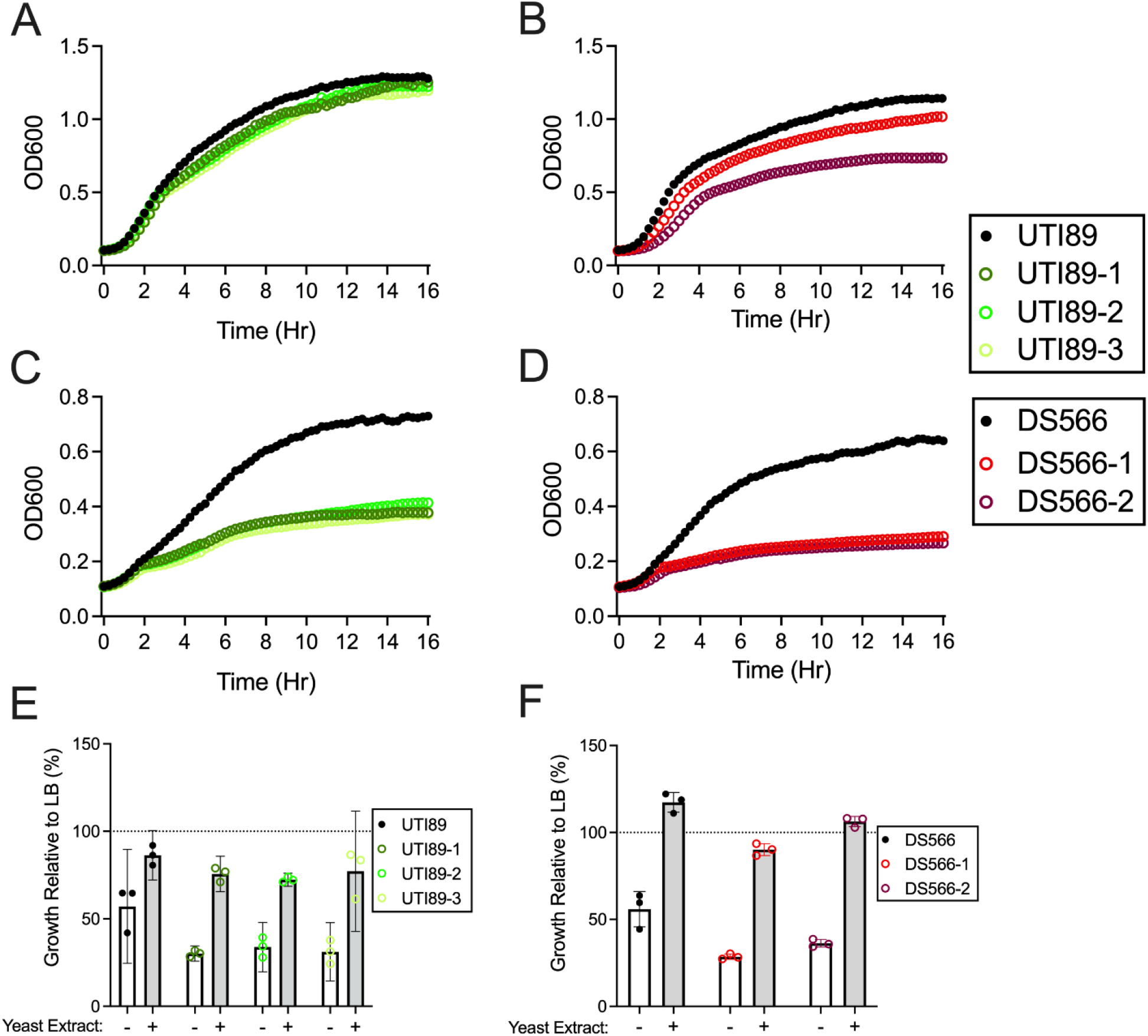
Phage resistant UPEC growth is attenuated in urine but can be partially restored with nutrient supplementation. (**A**) Mutants of UTI89 and (**B**) DS566 grow similar to their parental strain in LB media, but their growth is heavily attenuated in pooled human urine (**C, D** respectively). (**E**) UTI89 and (**F**) DS566 and their phage-resistant mutants were grown in urine with or without yeast extract supplementation or LB media 16h. Percent growth relative to LB for urine (white bar) and supplemented urine (grey bar) is displayed. Growth in LB (A,B) and urine (C,D) are representative of three independent experiments performed with four technical replicates. Growth in urine supplemented with yeast extract (E,F) is representative of three independent experiments performed in technical triplicate.

Defective growth in urine suggests that urine either contains active inhibitory factors for these phage-resistant bacteria or that LPS is important for growth in nutrient poor conditions. To delineate the reason for this growth defect, we supplemented urine with yeast extract, a primary nutrient source in LB medium. When bacteria were exposed to these conditions, their growth was partially rescued to levels resembling their growth in LB medium (**Fig. 5E-F**). This suggests that LPS is important for nutrient acquisition in nutrient-deplete conditions such as human urine.

### LPS mutations decrease murine bladder colonization

Because phage resistant mutants had comparable or increased biofilm formation, adherence, and internalization *in vitro*, we evaluated the ability of two LPS mutants (UTI89-2 and DS566-2) to colonize the mouse urinary tract compared to their parental strains. Bacteria were transurethrally instilled into the bladders of mice and, at 24 hours post-infection, bladder bacterial burdens were quantified. Wild-type UTI89 were capable of colonizing murine bladders while UTI89-2 was significantly attenuated in its colonization (**Fig. 6A**). While wild-type DS566 achieved lower bacterial burdens in the bladder than wild-type UTI89, LPS-mutant DS566-2 displayed a similar colonization defect, with bacterial burdens below the limit of detection in bladders at 24h post infection in all mice (**Fig. 6B**).

**Figure 6:**
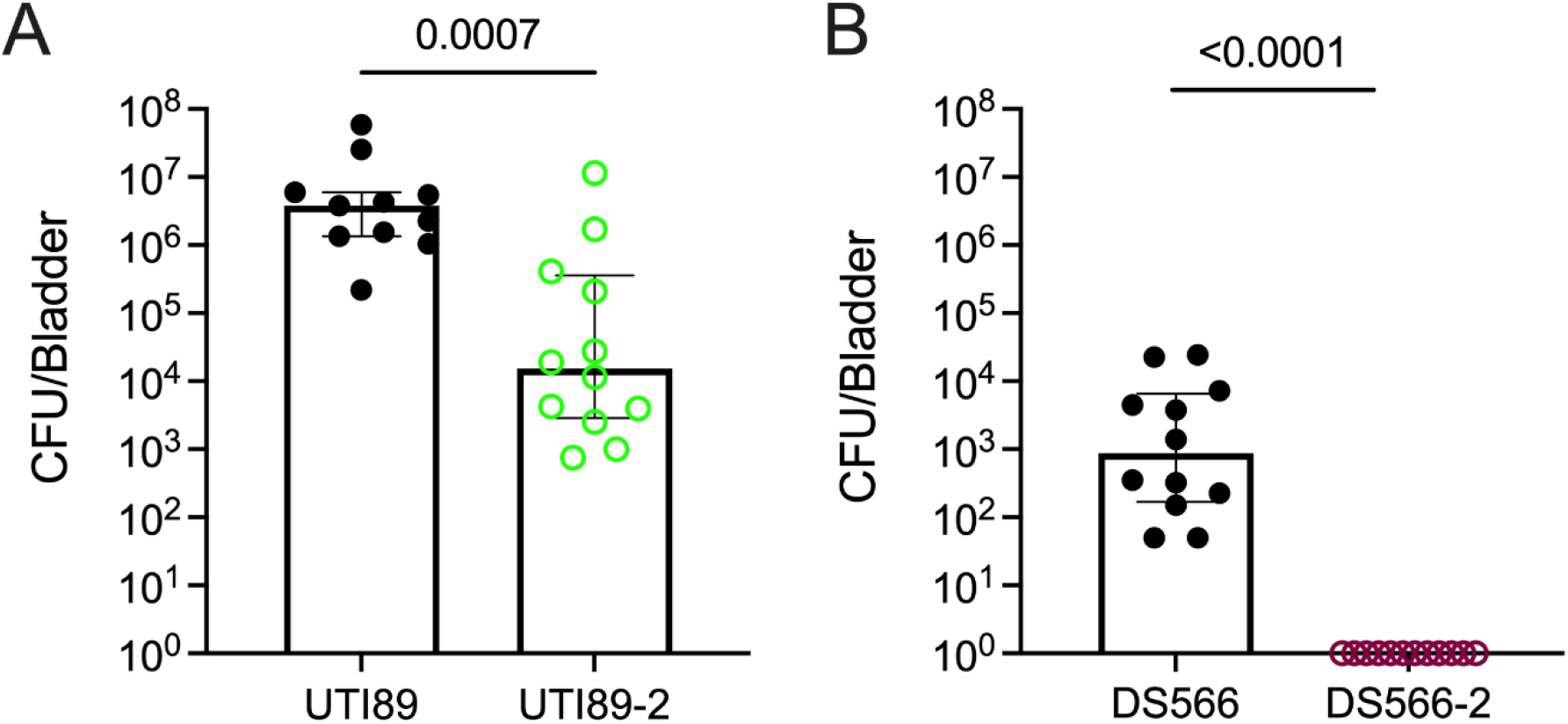
Phage resistant UPEC have decreased bladder colonization in a murine model of UTI. (**A**) Female C57BL/6J mice were transurethrally infected with 10^8^ CFU of UTI89 or UTI89-2 or (**B**) DS566 or DS566-2. After 24h bladders were removed, and bladder colonization was assessed. Points represent individual biological replicates performed over two separate experiments. (**A**) UTI89 *n* = 11, UTI89-2 *n* = 12; (**B**) DS566 *n* = 12, DS566-2 *n* = 12. Lines represent median and interquartile range. Data were analyzed by Mann-Whitney test.

## DISCUSSION

Although well-characterized in bacteriologic media, phage-bacterial dynamics are not well-defined in the urinary environment. Through our investigation, we identified (i) resistance arising to two genetically distinct phages, HP3 and ES17, in both bacteriologic medium and pooled human urine driven by mutations in LPS; (ii) that LPS truncation attenuates the growth of UPEC in urine, but that this phenotype can be partially rescued by supplementation with additional nutrients; (iii) that phage-resistant bacteria may be sensitized to membrane-interacting antibiotics; and finally, (iv) although LPS mutation may lead to increased adherence, invasion, and biofilm formation *in vitro*, this does not result in successful bladder colonization *in vivo*. In total, these finding suggest that phage-resistant bacteria may arise during phage therapy but that the resulting bacteria may be less capable of causing disease and may be sensitized to other treatment options.

Phage resistance is a well-documented phenomenon during bacterial infection with phage; however, few studies to date have assessed phage resistance outside of bacteriologic media and in the environment relevant to human medicine. Since human urine is a significantly more complex and dynamic than bacteriologic media, understanding phage-bacterial interactions in this environment is critical for assessing treatment efficacy. Our results demonstrate that UPEC killing by phages HP3 and ES17 is not determined solely by environmental conditions, but is highly influenced by the genetics of the bacterial host as revealed by differences among UPEC strains. In both environments, phage resistance developed in a short time frame. ExPEC have previously been shown to evade phage killing by HP3 via mutating LPS(35). Indeed, the HP3-resistant UPEC isolated in our current study harbors a mutation in transcription factor rfaH, which is involved in the assembly of the LPS(67). Interestingly, ES17-resistant UPEC also harbored mutations in LPS biosynthesis and assembly pathways. LPS is a major component of the Gram-negative outer membrane and provides stability to the outer membrane while anchoring other outer membrane proteins in place(68). Studies have previously shown that LPS mutation in *E. coli* attenuates several cellular processes, including cellular membrane integrity, resistance to antibiotics, and growth in low pH and high detergent conditions(65, 69-71). Here, we demonstrate that these same detrimental effects occur in the urinary environment.

The urinary tract is complex and dynamic with changing nutrient and solute concentrations throughout the day and from person to person(72-74). UPEC resistant to HP3 and ES17 isolated were equally capable of growth in LB as their parental strains but grew poorly in urine. This suggests that LPS mutation as a consequence of phage resistance may either lead to a sensitivity to host urinary conditions, or that LPS is important for effective growth in a nutrient-limited environment. When we introduced yeast extract into urine, bacterial growth was rescued to levels resembling bacterial growth in LB. This suggests that the growth defect observed in urine is not due to an existing antimicrobial factor but is instead primarily due to an inability of LPS-mutant UPEC to effectively acquire key nutrients in deplete conditions. Outer membrane proteins are needed in Gram negative bacteria to take up nutrients through the outer membrane. Several of these proteins require LPS for their assembly and insertion into the membrane(65, 68, 75). The decreased efficacy or number of these outer membrane proteins may account for the need to increase the nutrient availability to overcome growth defects, however, this topic is outside the scope of this study.

Increased biofilm formation has been described in *E. coli* harboring LPS “deep-rough” mutations *in vitro*(76). While biofilms *in vitro* may differ from those *in vivo*, since enhanced biofilms could lead to worse outcomes in patients treated with phages, we assessed if the phage-resistant UPEC displayed this this same increased biofilm formation. Partially aligning with previous literature, we observed *in vitro* biofilm increases in all three of our UTI89 strains containing *rfaH* mutations, although increased biofilm formation was not uniformly observed across all conditions. In previous literature, biofilm formation was increased in *E. coli* containing *rfaH* mutations when grown in M63B1 minimal media, although differing *E. coli* strains, biofilm generation, and enumeration methods were used(77). Additionally, we observed increased biofilm formation in both DS566 strains harboring LPS mutations. This increase appears to be most pronounced in bacteria DS566-2, which has the most severely attenuated LPS structure. Although *in vitro* biofilm assays may not fully reflect capacity to form biofilms *in vivo*, our results correlate with those observed by Nakao et al. who demonstrated enhanced biofilm production by *E. coli* defective in LPS heptose biosynthesis and found increased cell surface hydrophobicity relative to the parental strain(76). A limitation of our study is that all phage-resistant mutants identified were isolated *in vitro*, yet previous work by our lab suggests that resistance is likely to develop *in vivo* as well(35).

As well as adhering to implanted devices such as catheters, UPEC adhere to and invade the bladder epithelium and establish intracellular bacterial reservoirs capable of reinitiating infection after treatment. We observed minimal impact of LPS modification on adherence to and invasion of HTB-9 cells in the UTI89 background, although adherence was significantly increased in phage-resistant mutants in the DS566 background. These results echo work done by Nagy *et al*. who found no changes in adherence to HCT-8 or INT407 intestinal cells in ExPEC lacking *rfaH*(78). In contrast to the adherence results, two of three UTI89 mutants showed increased invasion of HTB-9 cells. Avian pathogenic *E. coli* lacking *rfaH* are more readily engulfed by chicken macrophages than the wild-type strain, however, these bacteria are poor at growing after being engulfed(79). The mechanism for this increased uptake was not further investigated but could explain the increased invasion of HTB-9 cells in our study.

Since our results suggested that LPS modification could alter host-pathogen interactions, we used a mouse model of UTI to investigate *in vivo* infection outcomes. We found that LPS mutants in both UTI89 and DS566 backgrounds were worse at colonizing the murine bladder than the wild-type strains. RfaH, a regulator of LPS, capsule, and alpha-hemolysin, is critical for bladder colonization as inactivation of *rfaH* dramatically lowers recovery of UPEC strain 536 from the urinary tract by 21 days after infection(80). Additionally, Aguiniga *et al*. used targeted gene deletions to identify LPS domains essential for colonization bladder using UPEC strain NU14(81). They showed that LPS-deficient UPEC were not only worse at colonizing murine bladders after 24 hours, but they were also less likely to be present in bladders two weeks post-infection, a measure of bacterial reservoir formation. While the LPS mutations assessed by this group are not identical to those observed in our study, the functional consequences (outer membrane truncation, inner membrane truncation, etc.) are likely similar. Although our study does not investigate the immune response to LPS-modified UPEC, others have observed the “rough” LPS phenotype in several asymptomatic bacteriuria strains, suggesting that these pathogens may be less virulent in humans(82). In fact, asymptomatic bacteriuria isolates with truncated LPS, among other virulence gene mutations, have been suggested as possible competitors which would be intentionally introduced in patients to prevent UPEC infection(83-86). In agreement with our study, bladder colonization using these UPEC was not always achieved despite frequent bacterial inoculations, suggesting a possible colonization defect of these strains(86).

Phage and antibiotics have the potential to work together synergistically, with one sensitizing bacteria to the other and vice versa(87). We wondered if LPS mutant UPEC may be more sensitive to antibiotics which target the bacterial membrane in urine, a condition which has not been investigated previously. Colistin and polymyxin B both act to permeabilize the bacterial outer membrane by interacting with the lipid A portion of LPS(88). Through MIC assays, we determined an increased sensitivity to these two antibiotics for most phage-resistant strains tested. As observed in the biofilm and adherence and invasion assays, this increase was most pronounced in DS566-2, the strain with the most significant predicted LPS truncation. Other groups have shown an increased resistance to colistin in Gram negative pathogens lacking the lipid A portion of LPS(89, 90). Our phage-driven mutants retain lipid A, explaining their sensitivity to colistin.

In summary, as is often the case with antibiotics, bacteria quickly become resistant to phage under physiologic conditions, an apparent threat to their use. However, as demonstrated here, mutations providing phage resistance may come at the cost of bacterial fitness and ultimately reduce pathogenesis. Many other phage targets exist in addition to LPS(38) raising the appealing possibility that combining phages with distinct targets could both reduce bacterial burden and attenuate bacterial virulence of emerging resistant bacteria. These findings expand our knowledge of phage resistance of UPEC in the context of the urinary tract and support their continued development as a target for controlling UTI.

## AUTHOR CONTRIBUTIONS

JZ, AM, and KP conceived and designed experiments. JZ, JC, SO, MB, MM, VME, EH and BS performed experiments. JZ, AM, and KP analyzed and interpreted results. BT provided clinical isolates and BT, AM, and KP acquired funding. JZ, AM, and KP drafted the manuscript. All authors contributed the discussion and manuscript edits.

## ACKNOWLEDGEMENTS

We are grateful to Dr. Sabrina Green for providing phage for experiments. JZ and MM were supported by an NIH T32 award (T32GM136554), and VME was supported by a scholarship from Baylor Research Advocates for Student Scientists (BRASS). Studies were supported by an NIH NIAID U19 award (U19 AI157981) to AM and KP. The funders had no role in study design, data collection and interpretation, or the decision to submit the work for publication.

