## Supplemental Material for "Phage resistance accompanies reduced fitness of uropathogenic *E. coli* in the urinary environment"

### Supplemental Tables:

| Strain | Type | Reference | Alternate | Strand | Nucleotide Position | Amino Acid Position | Effect | Gene | Product |
|---|---|---|---|---|---|---|---|---|---|
| UTI89-1 | Deletion | ACCAGGTG | A | - | 483/933 | 159/310 | Frameshift | tsx | nucleoside channel; receptor of phage T6 and colicin K |
| UTI89-3 | SNP | G | **A** | - | 82/936 | 28/311 | Stop Gained | wciP | putative rhamnosyltransferase |
| DS566-1 | SNP | A | C | - | 596/1569 | 199/522 | Missense | | cytochrome bd-I ubiquinol oxidase subunit 1 |
| DS566-1 | SNP | G | A | + | 478/795 | 160/264 | Missense | | hypothetical protein |

**Supplementary Table 1: Non-LPS mutations identified during sequencing.**

**Supplemental Figures:**

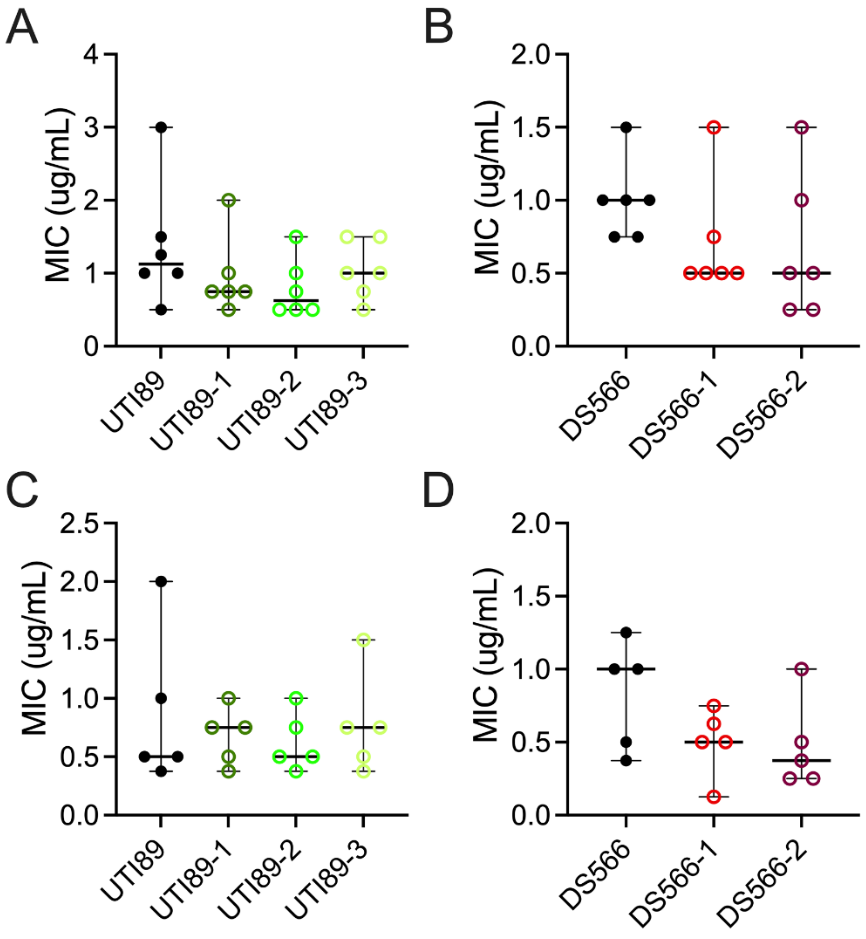


**Supplementary Figure 1: MICs for antibiotics targeting the bacterial outer membrane are not altered when LPS mutant bacteria are grown in LB media.** Colistin MICs of (**A**) UTI89 and (**B**) DS566 and their LPS mutants. Polymyxin B MICs of (**C**)UTI89 and (**D**) DS566 and their LPS mutants. Individual points are representative of independent experiments performed in duplicate. Bars represent median and 95% confidence intervals.
